# An Integrated Approach to Tracking Mandibular Position Relative to Incisal and Condylar Envelopes of Motion During Intraoral Clinical Procedures: A New Look at TMJ Movements

**DOI:** 10.1101/2025.03.11.642654

**Authors:** Jing-Sheng Li, Douglas S Ramsay, Andrea B. Burke, Greg J Huang, Fritzie I Arce-McShane

## Abstract

Current methods for tracking temporomandibular joint (TMJ) movements are difficult to perform during dental procedures. Yet precise and accurate quantification of mandibular movements is critical for understanding temporomandibular biomechanics and how it may contribute to temporomandibular dysfunction. This is particularly relevant to clinical procedures that might move the mandible near or beyond its functional range of motion. We present a novel approach that integrates cone-beam computed tomography, optical intra-oral scans, and six degree-of-freedom electromagnetic sensor data to quantify mandibular movements. This method employs rigid body transformations to generate subject-specific three-dimensional (3D) envelopes of motion and assess whether incisal and condylar landmarks remain within their functional 3D envelopes of motion. We demonstrate the clinical utility of this approach through simulated mandibular poses presented relative to the limits of incisal and condylar envelopes created from that individual’s voluntary border movements. Our findings reveal that condylar or incisal points in simulated mandibular poses are located beyond their normal motion envelopes, highlighting the importance of simultaneous monitoring of incisal and condylar landmarks. This methodology provides a clinically relevant tool for understanding temporomandibular biomechanics and it has the potential to signal clinicians when jaw movements during dental and oral surgical procedures approach or exceed the jaw’s functional range of motion and thereby could prevent adverse effects on the TMJ.

## Introduction

Temporomandibular disorder (TMD) is the second most common chronic musculoskeletal condition after chronic low back pain, affecting 5-12% of the population at a cost of approximately $4 billion annually [1]. TMD is an umbrella term that encompasses numerous clinical conditions involving the masticatory muscles, the temporomandibular joint (TMJ), and associated nerves and tissues [2]. With more than 30 TMDs in the expanded taxonomy [3], research investigating the diagnosis, etiology, prevention, and optimal treatment of TMDs is critical.

Injury serves as a triggering factor for TMDs [4–9] as evidenced by a prospective study [6] that revealed a 4-fold elevated risk of TMD onset following jaw injury. Ohrbach and Sharma [8] recently concluded that “Injury can qualify as a true etiology for TMDs”. The unexpected nature of injuries hinders a detailed clinical investigation of how the mandible and associated soft structures are stretched, compressed, or damaged. However, elective intraoral clinical procedures, such as third molar removal (3MR), would be amenable to detailed kinematic investigation. A systematic review [10] of 3MR and the subsequent development of signs and symptoms of TMD concluded that they are associated and influenced by factors such as: gender, age, third molar location, severity of impaction, and surgical difficulty. To explain this association, Huang and colleagues [5,11] hypothesized that during 3MR, the mandible is positioned in ways that could potentially harm the TMJ complex. These injuries may subsequently contribute to the development of orofacial pain/TMD symptoms. Testing this hypothesis requires measuring mandibular movements relative to the cranium intraoperatively during 3MR. Currently available mandibular tracking methods are not suitable for use during surgery other than surgical navigation system (e.g., large appliances placed intraorally, optical line of sight required throughout surgery, magnetic field source placement that interferes with surgical access) [12]. The goals of this project were to develop a method that could be used intraoperatively and to demonstrate its application.

The TMJ functions as a bilateral synovial joint, located where the mandible’s right and left condyles articulate with the articular fossa and articular eminence of the temporal bone. A fibrocartilaginous articular disc is positioned between the poorly congruent articulating surfaces of the condyle and the temporal bone. Discal ligaments attach the condyle to the disc, which allow the disc to move along with the condyle as it translates (glides) along the articular eminence while also limiting the condyle to rotational movements in the lower joint compartment [13]. Other ligaments are pertinent to TMJ function [14–16]. For example, the bilateral temporomandibular ligament (TML), which connects from the articular eminence at the articular tubercle to the condylar neck reinforces the lateral aspect of the capsule. The TML and capsule limit the condyle from being distracted from the articular eminence, and the TML would tense on retrusion, especially the one-sided retrusion that occurs with ipsilateral excursion. The stylomandibular ligament, which connects from styloid process of the temporal bone to the angle of the mandible, has been suggested to protect the TMJ by limiting mandibular protrusion [17]. The sphenomandibular ligament connects from the sphenoid spine to the lingula of the mandible and is not thought to play a significant role in mandibular movement, although it has been suggested to limit excessive translation of the condyle after 10° of opening [17].

The anatomical interplay between the mandibular fossa of the temporal bone and mandibular condyles, together with other tissues surrounding the TMJ, are crucial for a comprehensive understanding of TMJ biomechanics. The mandible can move in 6 degrees of freedom (*dof*) because the condyle and the temporal bone are regarded for kinematic purposes as separate rigid bodies in space. Although the mandible can move with 6dof, its position does have some constraints from surrounding structures, such as articular surfaces of the temporal bone, occlusion of upper and lower teeth, muscles, and ligaments. The mandible makes 2,000 or more movements per day [18]. Mandibular movements are commonly portrayed using an incisal envelope of movements, or Posselt’s envelope [19–21], that is generated using paths traced by a reproducible landmark often on a mandibular central incisor (e.g., medial incisal corner). The vast majority of movements fall within the envelope of border movements [22,23] such as those that occur during speaking, biting, chewing, and swallowing. The more extreme border movements of the mandible are thought to be constrained by anatomic limitations of the masticatory system that include skeletal, dental, ligamentous and muscular components as well as by proprioceptive and pain perception [24].

Various tracking systems have been developed to study the kinematics of the mandible, such as mechanical-linkage [25], optoelectronic [26–28], electromagnetic [23,29], radiographic[19], [20], or ultrasonic systems [25]. Early mechanical methods like pantographs and axiography provided limited spatial resolution. Radiographic methods offer detailed spatial resolution but expose patients to ionizing radiation. Ultrasonic systems are non-invasive and affordable but suffer from lower accuracy and are susceptible to ambient noise and reflections. A recent review [12] comparing these systems reports that ultrasonic tracking and axiography are fast and cost-effective but have higher measurement errors compared to optoelectronic and electromagnetic systems.

Incisal point-based kinematics has been widely used clinically due to the ease of tracking an incisal point and the simplicity of describing its kinematics, and it may provide clinical insights into normal and pathological conditions as they reflect the neuromuscular control of the mandibular motion [22,23]. However, tracking a single point to illustrate mandibular motion is problematic because it cannot describe the movements being made by the mandible given that a point can move with 3dof while the mandible can move with 6dof [32–34]. With the use of subject-specific morphology from medical imaging, such as computed tomography (CT), cone-beam computed tomography (CBCT), or magnetic resonance imaging (MRI), current optoelectronic and electromagnetic methods could describe the condylar movements [27,29,35,36] but have not been used extensively to track TMJ movements during dental procedures. The optoelectronic tracking system often mounts markers from the extension of the teeth that may directly interfere with performing intraoral clinical procedures. Also, it requires line-of-sight and can interfere with dental procedures or require clinicians to change their preferred posture due to cameras and markers that obstruct the operating field. In contrast, electromagnetic tracking systems [22,35,37] do not require line-of-sight, supporting its potential use during dental procedures.

Here, we present a workflow of tracking subject-specific mandibular movements during intraoral clinical procedures. The proposed method uses CBCT, optical intra-oral scans, electromagnetic 6dof sensors, and a small magnetic field source, to estimate 3D mandibular positions and orientations relative to the cranium. Also, we present case simulations of bilateral condylar position and orientation near or beyond an individual’s envelopes of motion. This approach will allow clinicians to gain a better understanding of the mandibular functional dynamics during treatment, and lead to better-informed decisions about the prevention, diagnosis, and treatment of TMD and other TMJ-related pathologies.

## Methods

The workflow starts from the creation of a custom retainer to house the microsensors for the mandible, followed by a case study to demonstrate how to collect a range of mandibular movements for generating incisal and condylar envelopes of movement during dynamic voluntary jaw movements. Four simulated poses demonstrate the interaction between the simulated incisal and condylar points.

### Creation of sensor-embedded retainer

This manufacturing process started with creating a fused cone-beam computed tomography (f-CBCT) to construct a precise 3D morphology of the patient’s dentition and craniofacial structures. A CBCT scan (3D Accuitomo 170, J. Morita, Saitama, Japan) is fused with an intra-oral scan (IOS, iTero Element Plus Imaging System, Align Technology, Tempe, Arizona, USA), to create a f-CBCT that provides a morphological description in sub-millimeter scale (**Fig. 1A**). The CBCT scan was taken with the upper and lower teeth together at the Maximal Intercuspation (MIP) position using a field of view of 140 × 100 mm with a voxel size of 0.270 mm. The CBCT images included the cranial and maxillary complex, TMJ, and mandible. Using parallel confocal imaging technology with optical and laser scanning, the IOS captured the high-accuracy surface geometry and color of the maxillary and mandibular teeth and soft tissues. The CBCT and the IOS images were uploaded to Relu (Leuven, Belgium). Relu automatically segmented the mandible from the craniomaxillary complex and individually segmented teeth from the IOS that were then superimposed on CBCT crowns to create an optimized hybrid image of optically scanned tooth crowns and gingiva from the IOS onto the CBCT to create a fused-CBCT (f-CBCT) (**Fig. 1B**). The gingiva on the mandible in the f-CBCT was omitted in our current application.

**Figure 1.**
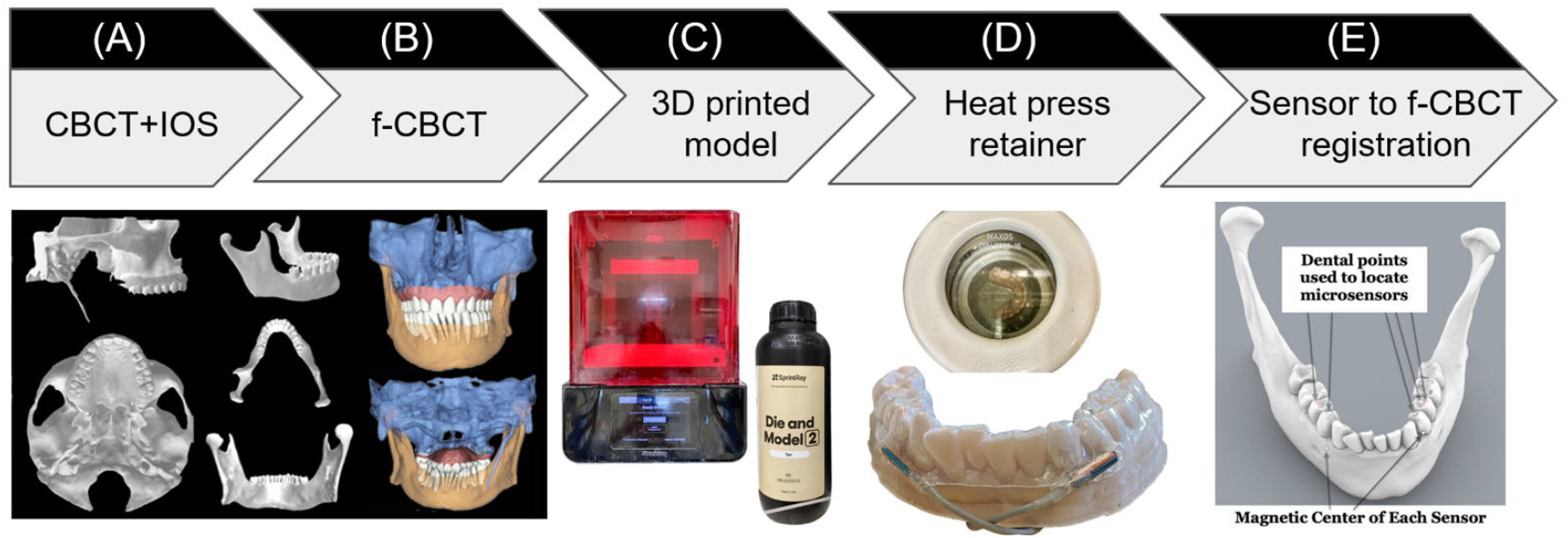
Sensor-embedded retainer for kinematic tracking. (**A**) Cone-beam computed tomography (CBCT) and intraoral scans (IOS). (**B**) CBCT and IOS were registered to create high-resolution fused (f-CBCT). (**C**) 3D printed f-CBCT model as a mold for retainer creation. (**D**) Use heat press to make sensor-embedded retainer. (**E**) Use dental points to locate microsensors in 3D space.

Maxillary and mandibular models were 3D printed (SprintRay, Los Angeles, CA) from the IOS scans. This 3D-printed mandibular model was used to fabricate the sensor-embedded retainer. Two “dummy” six-degrees-of-freedom (6dof) micro electromagnetic sensors (Micro Sensor 1.8, Polhemus, Colchester, VT) were placed buccal to the mandibular crowns of the canine and first premolar near the gingival margin on the right and left sides (**Fig. 1C**). The retainer was created using biocompatible thermoforming material (Zendura FLX, 0.76-mm thick, Bay Materials, Fremont, CA) with the heat pressure-form machine and trimmed to fit over the crowns of the upper and lower teeth. The dummy sensors were removed from the lower retainer and the actual sensors were inserted in their places. After the lower model had been coated with silicone separating medium to prevent adhesion, light cure aligner adhesive (Bond Aligner, Reliance Orthodontic Products, Itasca, IL) was applied around each microsensor in the retainer. The retainer was then fitted on the lower model’s teeth, and the adhesive was light-cured to fix the microsensors in the retainer (**Fig. 1D**). Precise 1-mm holes were placed through the retainer centered over each unique dental feature previously identified as visible on both models and on the f-CBCT. To determine the position of two microsensors with respect to the f-CBCT (**Figure 1E**), the subject wore the lower retainer and the tip of a hand-held 6dof stylus (3D Digitizer, Polhemus, Colchester, VT) was inserted into each hole in the retainer to touch the dental feature. The digitizing stylus located each previously defined dental features relative to the simultaneously located microsensors which were then registered through an interactive closet point method [25] on the f-CBCT. This allows 3D animation of mandibular movements tracked by the microsensors.

### Mandibular motion capture

A co-author participated as a subject (a healthy male, age 66, with no history of TMD symptoms related to jaw movements). We followed the workflow above to create a custom sensor-embedded retainer. Four 6dof sensors (RX2 Standard Sensor, Polhemus, Colchester, VT) were placed on the forehead to serve as reference points for jaw movements and the retainer with two microsensors was placed on the mandibular teeth.

While wearing the sensor-embedded mandibular retainer, the participant was instructed to perform a series of jaw movements in the sagittal, coronal, and transverse planes [39]. After a warm-up session, we recorded a dataset comprised of 5 repetitions of each of the following movement tasks: 1) Mouth opening from MIP position and closing, 2) Jaw moving laterally to the far right or left from MIP position, 3) Jaw moving forward or backward from MIP position, 4) combined movements: mouth opening at ∼6mm, 25%, 50%, 75% and 100% from MIP position, and then jaw moving laterally to far right or left, or starting at maximum protruded position, and jaw moving laterally to far right or left (**Fig. 2**). The sampling rate for these recordings was set at 240Hz to ensure high-resolution data capture. No hardware filtering was applied during data collection.

**Figure 2.**
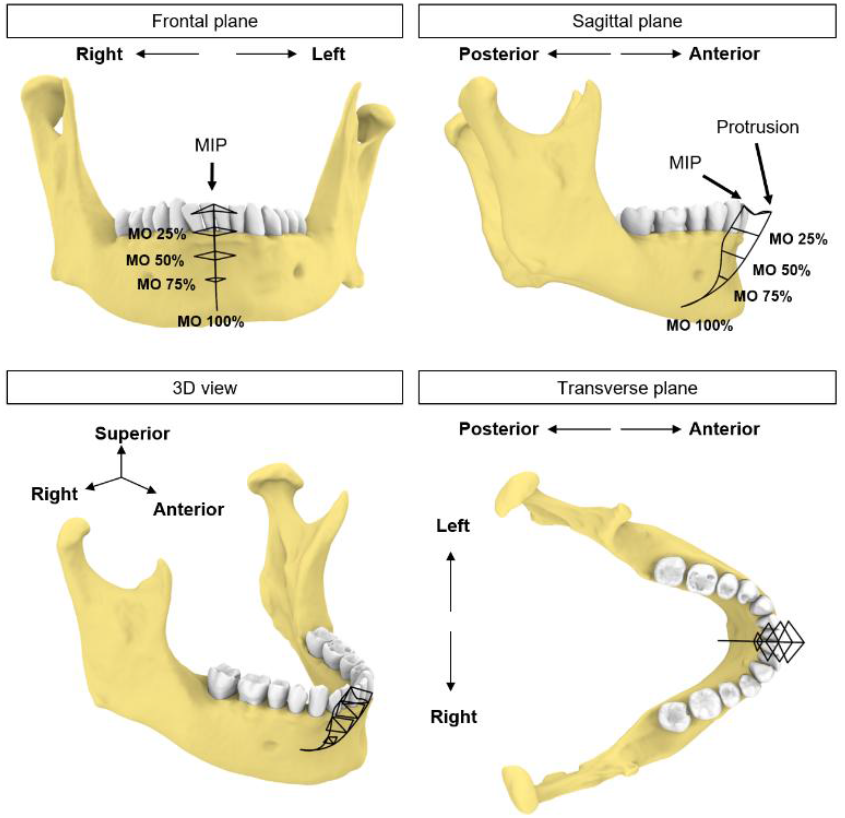
Schematic illustration of incisal trajectories (black solid lines). The participant uses this figure as a guide to move the jaw to different levels of mouth opening (MO) starting from MIP position or protruded position.

### Postprocessing

The first step is to orient the f-CBCT in a common frame and interpretable coordinate system. We describe the movement using the cranial coordinate system based on the maxillary occlusal plane, and anatomical alignments [40]. The XYZ axes correspond to movements along the medial-lateral (left-right), anterior-posterior (front-back), and superior-inferior (up-down) directions, respectively. The X-axis represents the anteroposterior direction, the Y-axis represents the transverse (buccolingual) direction, and the Z-axis represents the vertical direction. Specific definitions are in the Appendix.

To generate 3D envelopes of motion, we first identified the incisal and condylar points from the f-CBCT model. The ***incisal point*** was selected as the point on the lower left central incisal edge in the X-Z plane. The ***condylar points*** were defined as the most superior (distant) point of each condyle based on the length of a perpendicular line from the X-Y plane. For each dataset, we used the first frame of the four cranial sensor locations as a reference and then registered the rest of the frames to the reference points. We determined the start and end of each mandibular trajectory using 3D velocity thresholds, then normalized the trajectory to 100 percent.

Incisal point movements were reconstructed by applying rigid body transformation matrices formed by the microsensors with respect to the four cranial sensors. Condylar movements were reconstructed using the same transformation matrices. No additional filtering was performed for the incisal and condylar movements. Since all movements started from the MIP position, we combined all recorded movements and used the previously identified incisal and condylar points to construct the complete incisal and condylar envelopes of movements.

### Tracking performance assessment

To evaluate the tracking performance, we calculated the Euclidean distance between the two microsensors and cranial sensors in the following conditions:

1. To represent the overall tracking consistency, we estimated the variability of the Euclidean distance between the two microsensors during all voluntary movements.
2. To examine the effect of the metal surgical instruments interference, a metal tongue retractor was moved in and out of the participant’s mouth that was held open by a bite block. We calculated the Euclidean distance and ±1 standard deviation (SD) between two microsensors during a “quiet” phase without the tongue retractor. The standard deviation of the distance in the quiet phase was set as threshold for signal distortion. When the tongue retractor was inside the mouth, the Euclidean distance between each microsensor and each of the four cranial sensors was compared against the threshold values during the quiet phase.

### Simulation

In this section, we simulated four conditions to demonstrate the interplay between the incisal and condylar movements and the importance of simultaneous monitoring of incisal and bilateral condylar points. Simply relying on incisal point alone to represent mandible pose is insufficient and may allow condylar poses outside their envelopes. These simulated cases also tended to portray conditions during dental procedures. For example, sedated patients have the mandible positioned by the clinician while conscious patients are instructed to position their mandible, sometimes with assistance by the clinician, to create working space for the dental procedures, such as open their mouth wider or move jaw to left or right. In these simulations, we consider the lower teeth and mandible as a single rigid body based on the assumption that there is no relative movement between any teeth nor between the teeth and the mandible.

### Case 1 – excessive mouth opening

To replicate a pose that exceeds the subject’s maximal mouth opening, we simulated incisal point at 110% of mouth opening. This was calculated based on the incisal trajectories during mouth opening from the most protruded position (0%) to full (100%) mouth opening. For rigid body registration, the mandible was aligned to the simulated incisal point (110% of mouth opening), while keeping the two condylar points at 100% to avoid assumptions on their positions at 110% of mouth opening (**Fig. 6B**).

### Case 2 –mouth opening with condyles remaining in the fossa

During the early phase of mouth opening, the mandibular condyles remain within the mandibular fossa until the separation between the mandible and the anterior teeth reaches 20 to 25 mm. Beyond this point, the mandible begins to translate out of the fossa [39]. To provide adequate access for instruments and proper visualization of the back teeth procedures, clinicians typically open the mouth of the patient (under sedation or general anesthesia) to about 40 mm while ensuring the condyles remain in the fossa by not allowing anterior translation [41]. Based on these data, we simulated a pose where the condylar points were only rotated along its axis, without translation, until the mouth reached a 40 mm linear distance from the incisal point at MIP (**Fig. 7B**).

### Case 3 – excessive lateral excursion of the jaw

In this simulation, we replicated a pose to expose the upper right molars by shifting an incisor 3mm beyond the rightmost border of the motion envelope at 50% mouth opening. The condylar points were aligned to the simulated incisal point (**Fig. 8B**).

### Case 4 – lateral excursion of the jaw with greater interocclusal space on the same side as the excursion

This simulation created additional vertical space for the right molar by lateral excursion of the jaw to the right. With the incisal point at 50% mouth opening and at the rightmost position of the envelope, we simulated a 5° rotation along the axis formed by the incisal and left condylar point to increase vertical space for the right molar (**Fig. 9A**).

## Results

A total of 35 distinct movement tasks were performed, and 175 trials of these movements were analyzed to create the incisal and condylar envelopes of movement.

### Incisal envelope of movements

The incisal envelope of movements is characterized by a feather-like shape when trajectories of all instructed mandibular movements were plotted together (**Fig. 3)**. As can be viewed from the frontal and transverse planes (**Fig. 3A-C**), this subject made fewer lateral movements to the right. By creating a bounding box [42], we measured the dimensions of the envelope of motion for this subject as 29.0 mm, 47.8 mm, and 49.0 mm along the medial-lateral (ML), anterior-posterior (AP), and superior-inferior (SI) direction, respectively (**Fig. 3D**). Comparison of these dimensions revealed that this participant exhibited an asymmetric incisal envelope that slightly deviated to the right during mouth opening; at full mouth opening, the incisal point is approximately 6.3 mm to the right from the MIP (see frontal plane, (**Fig. 3A**). Moreover, the envelope became very narrow at nearly two-thirds of the mouth opening path (**Fig. 3B**).

**Figure 3.**
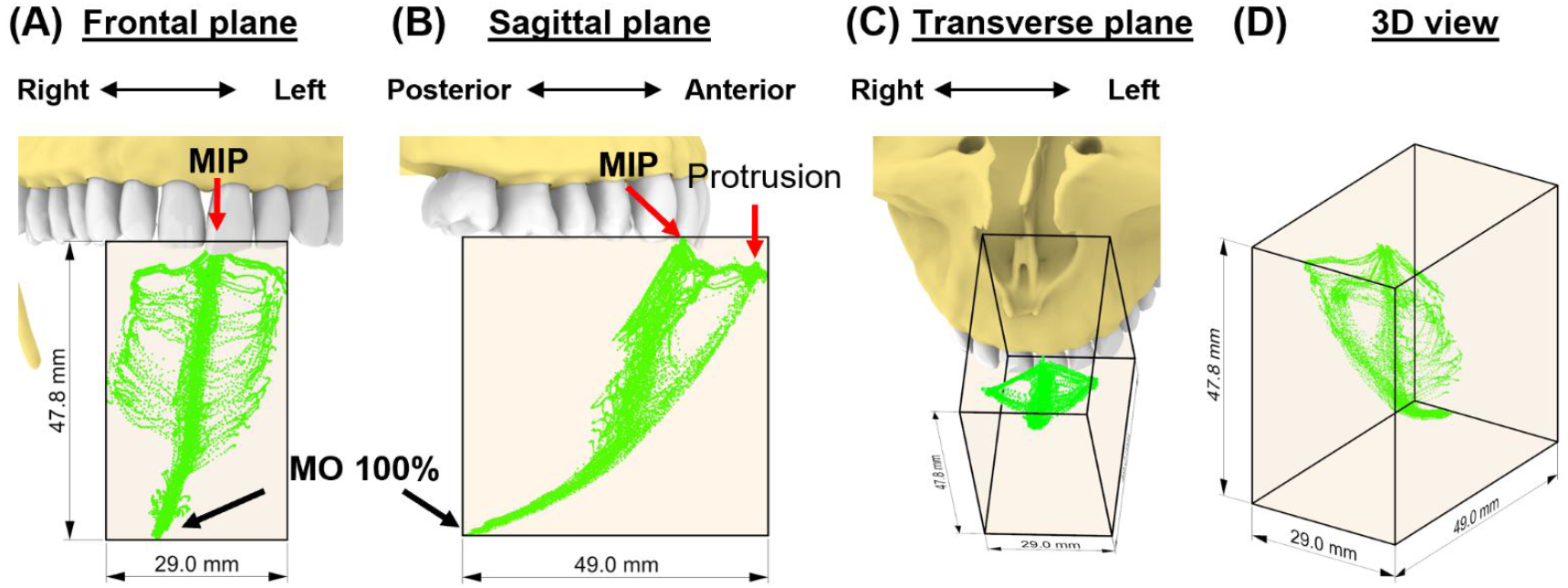
The subject-specific incisal envelope of movements is shown in frontal plane (**A**), sagittal plane (**B**), transverse plane (**C**), and in 3D (**D**) relative to the incisors. A bounding box for the incisal envelope was created to measure the dimension of the envelope.

### Condylar envelopes

The 35 movement tasks formed dissimilar shapes of envelopes for each condyle in this subject (**Fig. 4**). Specifically, the left condylar envelope was a J-shape while the right was a U-shape (**Fig. 4D-E)**.

**Figure 4.**
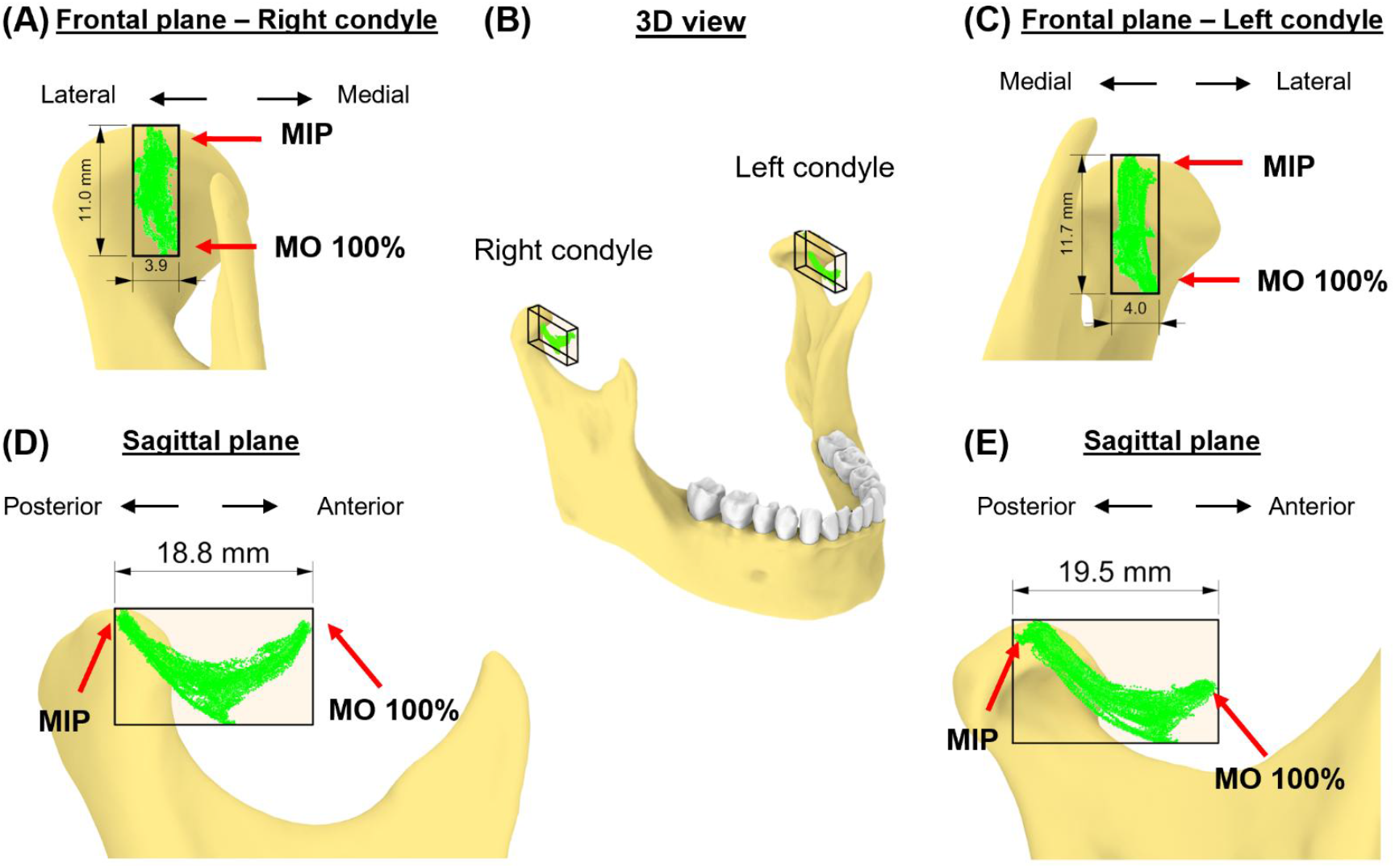
The subject-specific condylar envelope of movements is shown in frontal plane (**A and C**), in 3D (**B**), and sagittal plane (**D and E**) relative to the high-point of the condyle. A bounding box for the condylar envelope was created to measure the dimension of the envelope.

Both right and left condylar points moved inferiorly and anteriorly in the first half of the condylar path during mouth opening. The right condylar point moved anteriorly and upwardly in the second half (∼50% of mouth opening); the left condylar point moved anteriorly. At full mouth opening, the right condylar point was 9 mm upwards compared to the left side of 5 mm. These observations indicate asymmetrical movements between the condyles. Dimension-wise, the differences between the two condylar envelopes were within 1mm (**Table 1**).

**Table 1.** Dimensions of the condylar envelopes of movements.

| *Table 1. Dimensions of the condylar envelopes of movements.* |  |  |
| --- | --- | --- |
|  | Left condyle (mm) | Right condyle (mm) |
| Medial-lateral | 4.0 | 3.9 |
| Anterior-posterior | 19.5 | 18.8 |
| Superior-inferior | 11.7 | 11.0 |

### Consistency assessment

Over the 35 voluntary movement tasks (average recording duration: 28.9 (SD: 6.8) seconds), the standard deviation of the distances between the two microsensors was 0.148 (SD: 0.062) mm (range: 0.040-0.292). In the surgical instrument metal artifact test, a threshold of 0.042 mm distance between positions recorded by microsensors to determine periods of signal distortion (**Fig. 5A**). There were three outcomes: 1) both microsensors were not affected, i.e., the distance between each microsensor and cranial sensors remained within the defined threshold, 2) one of the microsensors had distorted signal, and 3) both microsensors were affected (**Fig. 5B**).

**Figure 5.**
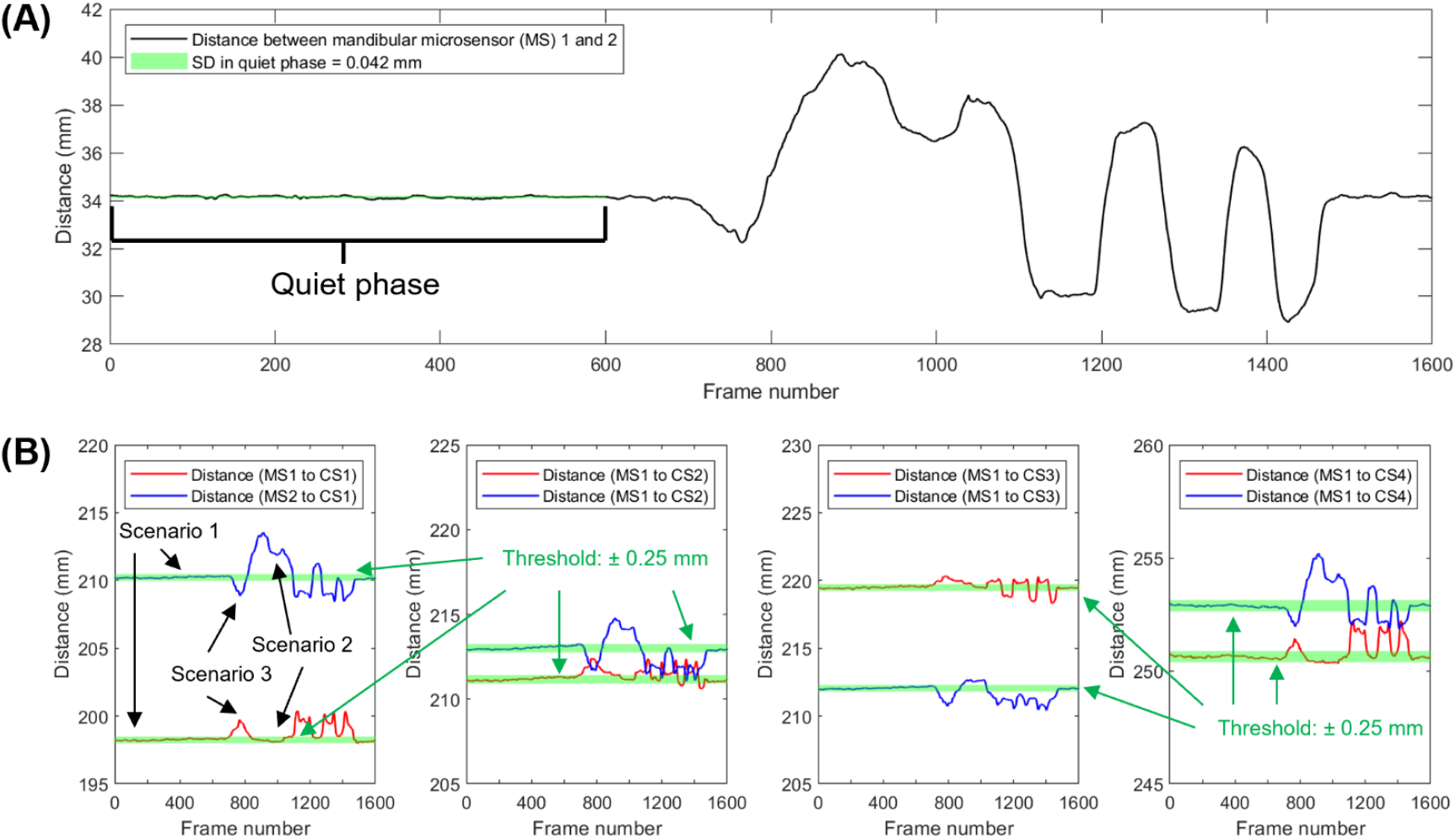
Approach used to identify sensor signal distortion. **(A)** Distance between mandibular sensors over time while a metal tongue retractor was placed and removed. (**B**) Distance between a specific cranial sensor (CS) and the left and right mandibular microsensors (MS). Scenario 1 shows a period of reliable signal from frames from 1 to 600, around frames 900-1500, deflections occur which indicate the presence of artifacts. Portions of the data show distortion of only one microsensor (scenario 2) or both microsensors (scenario 3).

### Simulations

In the first simulation (**Fig. 6**), we assessed the effects of an additional 10% mouth opening beyond the subject’s maximal voluntary opening. At the maximal protruded position, the mandible rotated 48.4° at 100% mouth opening along a circular path and 53.2° at 110%. At this simulated pose, both left and right condyles were 0.2 and 0.8 mm outside their condylar envelopes (**Fig. 6A-C**). The simulated incisal point was also 6.8 mm outside the incisal envelope (**Fig. 6D-E**). Thus, at 110% mouth opening, the incisal point exhibited the largest deviation from its envelope of motion.

**Figure 6.**
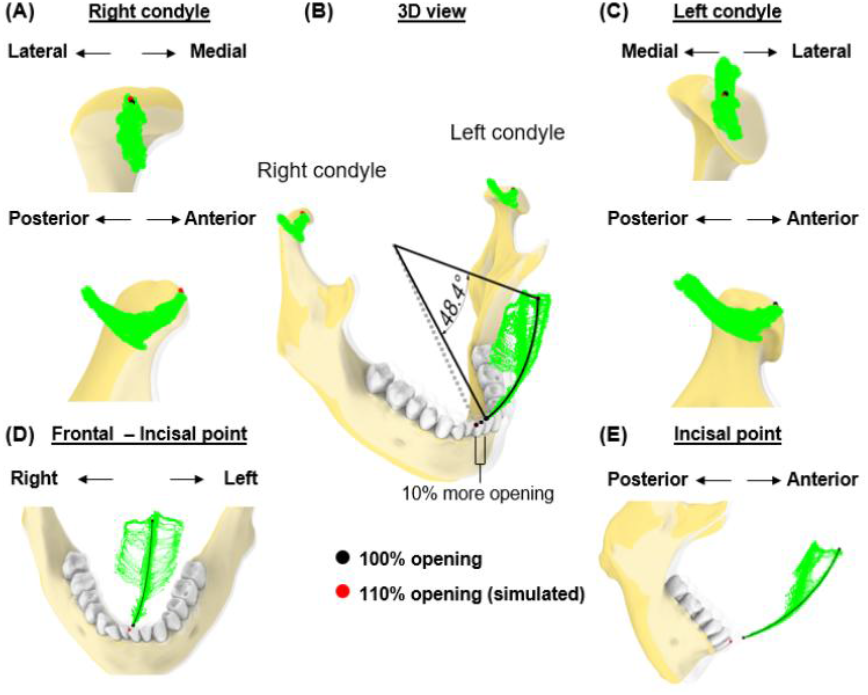
Simulation case 1. (**A**) Right condylar points at maximum opening (black) and simulated pose (red) are shown relative to condylar envelope. (**B**) Incisal and condylar points relative to their envelopes, shown at full mouth opening (transparent jaw) vs. 110% of mouth opening along a circular path. (**C**), As in (A), but shown for left condyle. (**D**-**E**) Frontal and sagittal views show incisal points relative to its envelope.

In the second case, we simulated a pose corresponding to a mouth opening of 40 mm (Euclidean distance) from the incisal point at MIP (**Fig. 7**). Mandibular rotation at this simulated pose was 24.1°, placing the incisal point 9.7 mm outside its envelope of motion.

**Figure 7.**
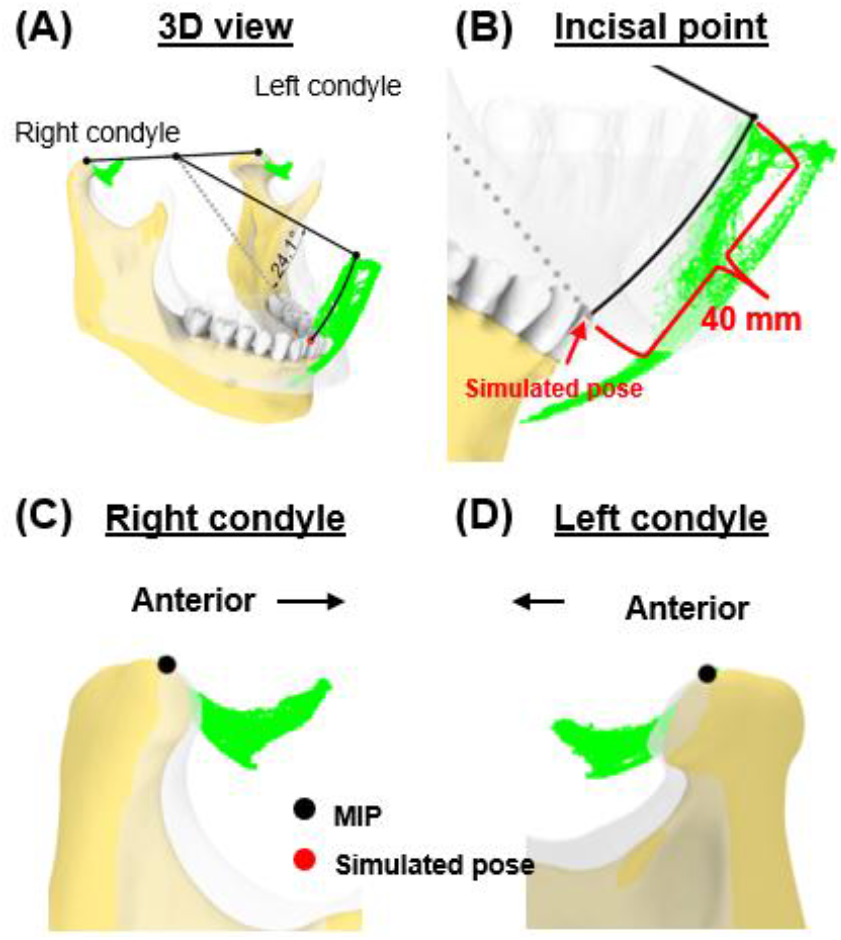
Simulation case 2. (**A**) Two condylar points are used to create a rotational axis used to simulate a mouth opening of 40mm. (**B**) Simulated incisal point is shown outside the incisal envelope of movement. (**C**-**D**), Condylar points at simulated pose remained unchanged.

Next, we simulated a jaw position that was shifted laterally (**Fig. 8**). At 50% mouth opening, we shifted the incisal point 3 mm from the rightmost border of the incisal envelope. In this simulated pose, the left condylar point was 2.1 mm outside the envelope while the right condylar point was outside its envelope by 0.1 mm (**Fig. 8A-C**). This indicates that excessive lateral movements (**Fig. 8D-E**) may lead to significant deviations from condylar envelopes, particularly affecting the contralateral condyle.

**Figure 8.**
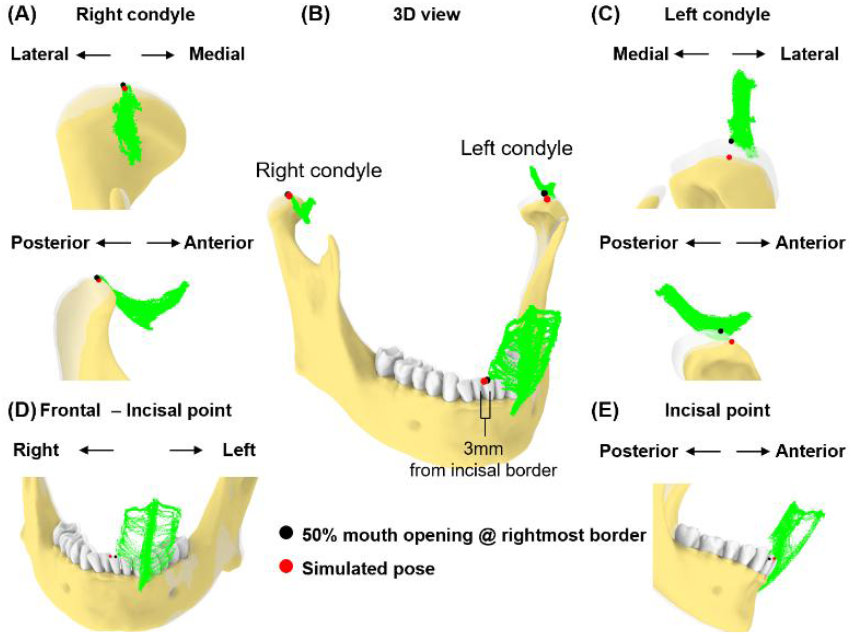
Simulation case 3. (**A**) Right condyle positions at 50% mouth opening to the rightmost border and simulated pose are shown relative to condylar envelope. (**B**) Illustration of a 3-mm incisal displacement to the right of the border of incisal envelope. Incisal and condylar points relative to their envelopes, shown at 50% opening with displacement to the rightmost border (transparent jaw) and simulated pose. (**C**) As in (A), but shown for left condyle. (**D**-**E**) Frontal and sagittal views show incisal point at 50% opening with displacement to the rightmost of the border and simulated pose relative to its envelope.

Lastly, we simulated a vertical space creation for back tooth procedures. The 5° rotation around the axis connecting the incisor and left condyle (**Fig. 9A-B**) resulted in 7.4 mm linear displacement from the original pose at the right condylar point (**Fig. 9C**). This places the right condyle outside of its envelope. Additionally, the linear distance between the upper and lower 2nd molars increased by 13.0% (from 21.6 mm to 24.4mm, **Fig. 9D**).

**Figure 9.**
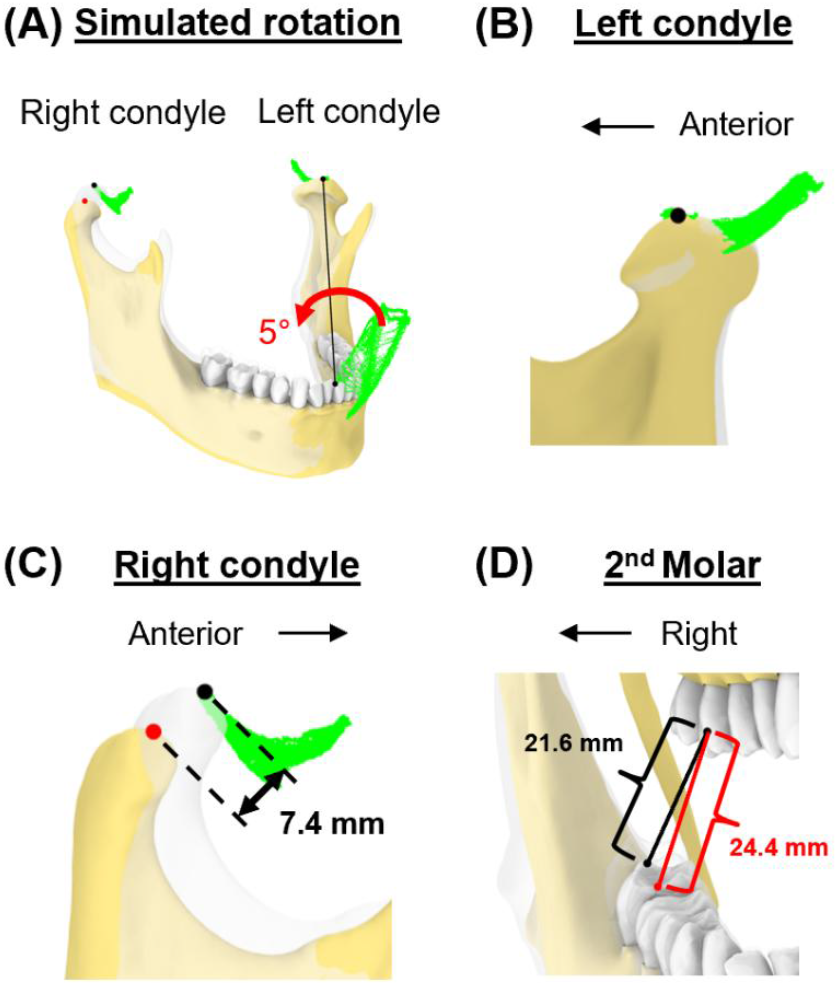
Simulation case 4. (**A**) Mandible is rotated for 5° around the axis connecting incisor and left condyle. (**B**) Left condylar points remain unchanged. (**C**) In this simulated pose, the right condyle was displaced by 7.4 mm. (**D**) The distance between the upper and lower second molars increased.

## Discussion

In this study, we introduced a novel approach for quantifying subject-specific mandibular movement dynamics. By simultaneously monitoring incisal and bilateral condylar envelopes of motion, our approach overcomes the inherent limitations of tracking incisal envelope alone, providing a more comprehensive and accurate 3D representation of mandibular dynamics compared to methods that rely solely on either incisal or condylar path analysis.

Several established methods have been developed to understand the basic functions of mandibular movements. For example, cine CT and cine MRI offers direct visualization of TMJ structures dynamically [31,43,44]. However, these measurements are not always feasible due to space limitations and practical challenges during dental procedures. Optical marker tracking system can adequately track voluntary mandibular movements by using external markers extended from the lower teeth [26,44–46]. Since optical marker tracking relies on a clear line of sight, any obstruction can cause tracking loss or errors, necessitating specific clearance for marker visibility during dental procedures.

In contrast, the electromagnetic tracking system used in our study eliminates the need for this clearance while providing accuracy comparable to the optical tracking system. However, the electromagnetic tracking system is susceptible to data distortion from metallic artifacts, such as when using metallic tools in the region between the sensor and magnetic source. This can be circumvented by using two microsensors to allow for data recovery in case of missing or distorted data from one of the microsensors. While both microsensors could be affected at the same time, we have presented a postprocessing method to distinguish and remove the distorted data. These results support the potential applicability of this electromagnetic tracking device for accurately tracking jaw movements during clinical procedures, allowing real-time feedback to clinicians. This is particularly significant in prolonged dental procedures, such as root canal therapy or the extraction of difficult-to-access impacted third molars on a sedated patient, which are linked to increased TMD prevalence or pain [47–49].

Our novel approach allows for the comparison of subject-specific incisal and condylar envelopes before and after clinical procedures. Combined with spatial and temporal information on mandibular positions during procedures, this information could provide new insights into TMD risk factors. Goldman [9] describes discal attachment injury as occurring when the mandible’s range of movement is exceeded or violated and results in tearing or loosening of the lateral discal and/or capsular ligaments. These discal ligaments are well vascularized and innervated and can become inflamed and painful [9] and what began with inflammation can lead to capsulitis. Injuries could occur, if a sedated patient’s mandible is moved passively near or beyond its borders, defined by awakened voluntary movements, that leads to ligaments and soft tissues being stretched or injured in positions that relate to rotational and translational movements of the mandible that are not obvious by examining a single incisal or condylar point relative to it its typical range of motion envelope.

The incisal envelope of movements generated in this study extends past research, which largely concentrated on sagittal plane movements, such as mouth opening and closing, or frontal plane side-to-side excursions. Past research which concentrated on sagittal plane movements has shown that incisal linear distance could be under 40 mm for patients with degenerative TMJ disorder [35], an average of low to mid-50 mm for adults [50,51], and an as high as 65.5 mm for orthodontic patients with different skeletal patterns [23]. Importantly, about half of the sample in the studies have a mean maximal opening smaller than the average value. The transverse plane at different levels of opening and combined-plane movements have not been studied explicitly [44]. The movement tasks outlined in this study could enable a more thorough construction of the “envelope of motion,” capturing both incisal and condylar movements completely.

There is a shape difference but slight variation in terms of dimensions between the two condylar envelopes of movement in our case study. The AP dimension in both condyles is less than 20 mm, which is consistent with the values reported in earlier research. Baltali and colleagues [35] reported AP condylar path of approximately 2 cm during mouth opening and closing for patients with degenerative TMJ disorder. Salaorni and Palla [51] reported slightly smaller condylar translations during mouth opening and closing (14 ± 4.2mm). Yatabe and colleagues [52,53] used OKAS-3D recording system, which does not include subject-specific imaging-based geometry, and reported the length of condylar movements in protrusive-retrusive movement and mouth opening-closing for approximately 23 mm for both condyles (SD: ∼4 mm). Huang and colleagues [44] reported approximately 19 mm (SD: 4mm) mouth opening and smaller movements during protrusion. Our study also demonstrates the presence of medio-lateral movements in the TMJ, although the range in this dimension is smaller compared to other dimensions, such as AP and SI movements. This finding is consistent with a study using single fluoroscopic imaging [30].

The incorporation of incisal and condylar tracking extends beyond jaw movement monitoring. The generated data allows for simulations that are useful for dental clinical practice, such as surgical planning and evaluation of potential risks of procedures to patients, and even as an aid for physical therapy of TMJ pain or dysfunctions. Our simulated cases demonstrated that excessive mouth opening or lateral excursion could result in asymmetric condylar displacement. This implies that conditions that cause excessive jaw translation and rotation may result in traumatic injuries to tissues around the condyles, such as the discal and sphenomandibular ligaments. Similarly, the elongation pattern of sphenomandibular or stylomandibular ligaments can be simulated to predict the impact of certain intraoral procedures. Understanding the interaction between ligament status and duration of loading can further enhance our knowledge of TMJ function and its implications for dental care.

## Conclusion

Our novel approach to quantifying subject-specific, dynamic incisal and condylar movements represents a significant advancement in TMJ biomechanics. By simultaneously tracking both incisal and condylar motion envelopes, we provide novel insight into the complex three-dimensional kinematics of mandibular function. This integrated perspective reveals critical relationships that remain invisible to traditional single-landmark tracking methods. While our findings demonstrate potential for clinical application, further validation across diverse patient populations is essential to refine this methodology and establish normative parameters. The next critical phase of research must focus on correlating these biomechanical measurements with clinical outcomes and patient-reported symptoms. By integrating our understanding of TMJ kinematics into clinical practice, we can potentially prevent damage to TMJ structures, develop targeted interventions for TMD, establish evidence-based parameters for safe mandibular manipulation during dental procedures, and improve individual patient outcomes.

## Author Contributions

**Jing-Sheng Li:** Conceptualization, Methodology, Software, Formal analysis, Investigation, Writing - Original Draft, Writing - Review & Editing, Visualization

**Douglas Ramsay:** Conceptualization, Methodology, Validation, Formal analysis, Investigation, Writing - Review & Editing, Supervision, Project administration, Funding acquisition

**Andrea Burke:** Conceptualization, Investigation, Writing - Review & Editing, Supervision, Funding acquisition

**Greg Huang:** Conceptualization, Investigation, Writing - Review & Editing, Supervision, Funding acquisition

**Fritzie Arce-McShane:** Conceptualization, Methodology, Validation, Formal analysis, Investigation, Writing - Review & Editing, Supervision, Project administration, Funding acquisition

