## Appendix for "An Integrated Approach to Tracking Mandibular Position Relative to Incisal and Condylar Envelopes of Motion During Intraoral Clinical Procedures: A New Look at TMJ Movements"

In this appendix, we provided the details of the coordinate system used in this study.

First, a mid-palatal raphe plane was defined by fitting a plane to the mid-palatal raphe points identified on the f-CBCT.

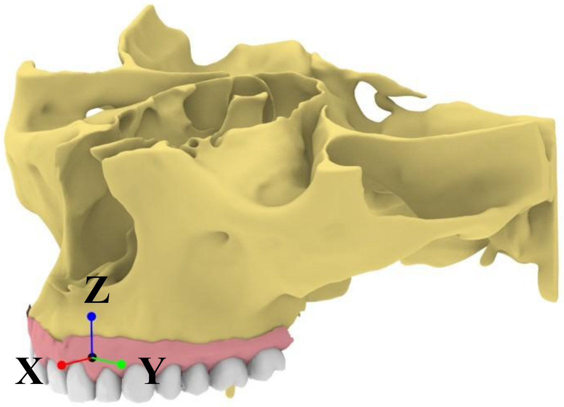


Next, an occlusal plane was determined by identifying the lingual supporting cusps of the maxillary posterior teeth that were in occlusal contact, as indicated by the IOS heat map.

We then drew a line at the intersection of the posterior region of the occlusal plane with the mid-palatal raphe plane; this line defines the anterior–posterior (AP) direction, oriented anatomically anteriorly.

To establish the coordinate axes, we took the cross product of (1) the normal vector to the mid-palatal raphe plane and (2) the unit vector corresponding to the AP direction. The resulting perpendicular vector defines the Z-direction (anatomically superior).

The origin (0,0,0) of this coordinate system was set at the most posterior aspect of the incisive papilla. Finally, the X-, Y-, and Z-axes were oriented to point anteriorly (X), to the left (Y), and superiorly (Z), respectively.
